# Seasonal Change of Microbial Diversity and Its Relation with Soil Chemical Properties in Orchard

**DOI:** 10.1101/600668

**Authors:** Xuhui Luo, Mingkuang Wang, Guiping Hu, Boqi Weng

## Abstract

This study aimed to determine the microbial diversity of different soil depths (0-5 and 5-20 cm) in a subtropical orchard during different seasons (i.e., Spring, Summer and Autumn) for enrich the knowledgements on micorbes roles in orchard ecosystem balance. In tracking experiments conducted in an orchard (established in 1996), the phospholipid fatty acid (PLFA) biomarker method was employed to know soil microbial system. Total PLFAs concentration did not vary significantly between soil depths but changed between seasons. It peaked in the summer at 258.97 ± 23.48 μg g^-1^ soil from 0-5 cm and at 270.99 ± 58.94 μg g^-1^ soil from 5-20 cm. A total of 33 microbial fatty acid biomarkers were observed and identified in the sampled soil. Quantities of PLFAs for 29 microbe groups varied significantly between seasons, except for 15:0 iso 3OH, 15:1 iso G, 16:0 2OH, and 17:0 iso 3OH. The bacterial PLFAs and fungal and actinomycotic PLFAs in the orchard soil collected in Summer were significantly higher than in the Spring or Autumn (*P* < 0.01). The number of soil microorganism species (Richness) and the Simpson and Shannon-Wiener indexes were all the highest in summer. The total PLFAs, bacterial PLFAs, fungal PLFAs, actinomycotic PLFAs, Richness, or the Simpson and Shannon-Wiener indexes were all significantly negatively correlated with soil pH, total carbon (TOC), total nitrogen (TN) and cation-exchange capacity (CEC) (*P* < 0.05).

## Introduction

The orchard ecosystem is important for fruit production and carbon sequestration, biodiversity decreases [1] and soil erosion [2], and pollution [3,4]. Hence, maintaining a balanced orchard ecosystem is essential. Soil microbes are essential for the functioning of terrestrial ecosystems because they play a unique and indispensable role in ecosystem balance [5].

Soil microbial community change by soil quality evolution because of long-term management. The interaction between microbes and soil quality is very complex. It is reflected in effects on microbial diversity of chemical properties change [6–11]. Spatial and temporal distribution of microbial community have some value information to explain the interaction [12–18]. However, few clear information had be found by observation on spatial and temporal distribution of microbial community in subtropical orchard.

Thus, we hypothesized that height seasonal temperature and moisture variation and vertical soil chemical properties change would lead to obvious differences in spatial and temporal distribution of microbial community in subtropical orchard. To demonstrate seasonal and vertical changes in microbial diversity and the links between soil microbial diversity and soil chemical properties in a subtropical orchard, PLFA was employed to monitor microorganism quantity and diversity in orchard soils under different seasons in the hilly red soil of the subtropical zone in southern China. The observations of this study regarding ecological parameters of orchard soil microorganisms can provide a scientific basis for further studies and management strategies.

## Materials and methods

### 2.1. Experiment Area

The experimental area was located at the Yuchi Village Experimental Station, Xicheng Township, Youxi County, Fujian Province, southeast China (26° 25’ N, 117°57’ E). The area has a subtropical humid monsoon climate with an actual annual sunshine time of 1781.7 h (accounting for 40% of the available annual sunshine hours); an annual precipitation of 1,284 mm (Fig. 1); an annual average temperature of 19.2°C; an average temperature in July of 26.6°-28.9°C and in January of 8.0°-12.0°C; and a frost-free period of more than 312 d. As an peach orchard water and soil conservation monitoring system, the experimental station was established in 1996 with an altitude of 150 m and a slope of 15° facing south-southeast. The soil was a Quaternary age red soil with a clay soil texture [19]. The experimental field was originally the secondary shrub-barren hill, and the constructive plant species were *Dicranopteris dichotoma* Bernh, *Miscanthus floridulus* War ex Schum, and *M. sinensis* Anderss. The surrounding vegetation was mainly coniferous and coniferous-deciduous-bamboo mixed stands with constructive species of *Pinus massoniana* Lamb and *Cunninghamia lanceolata* Hook and a grove of *Phyllostachys heterocycla* cv. Pubescens. The station was initially established for fixed soil erosion monitoring.

**Fig. 1.**
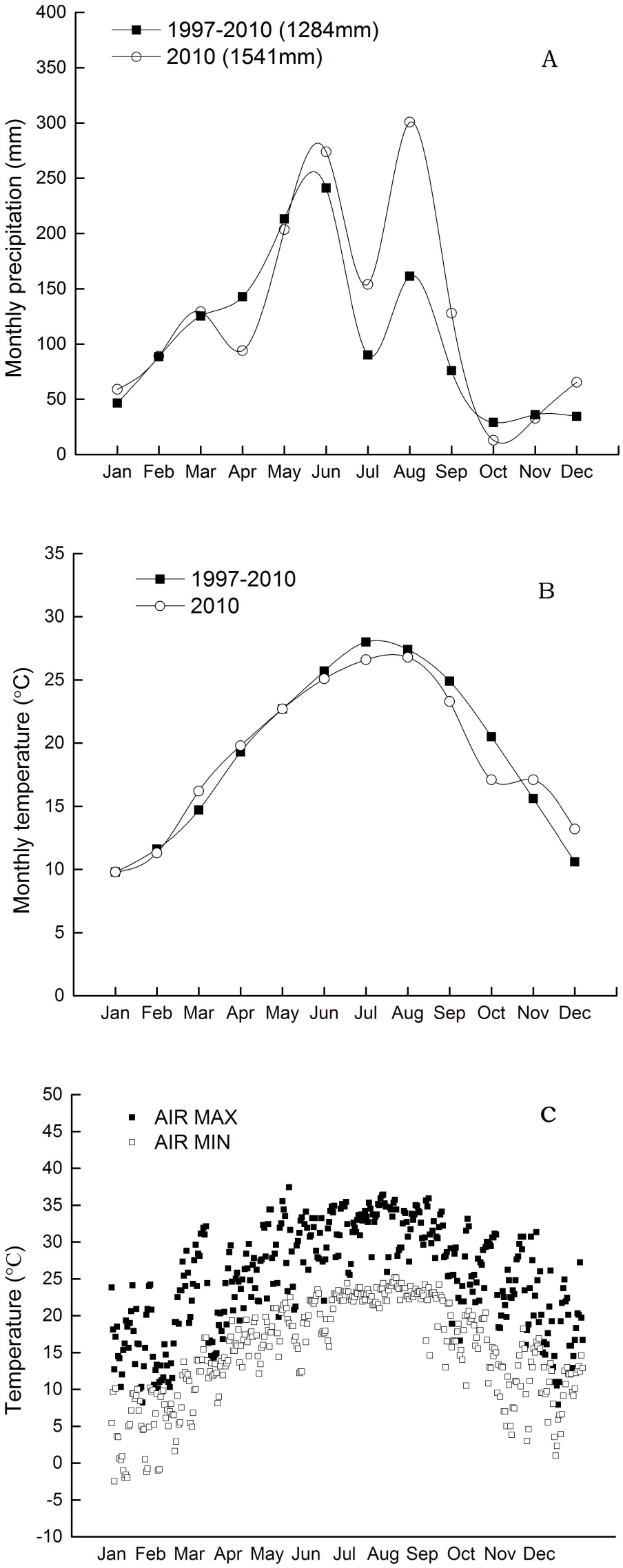
Monthly dynamics of precipitation (A), average air temperature (B), and extremes air temperature (C) at the trail location from 1997 to 2010. Annul precipitation is shown in parentheses (A).

### 2.2. Soil sampling

Spring, Summer and Autumn as main treatments had been settled for seasonal factors. On April 27, August 22, and November 4, 2010, at 3 days (d) after raining to keep soil moisture at the same level, soils were sampled from three random spots in the non-fertilizing zones of the area surrounding the average tree were collected from the soil depths of 0-5 and 5-20 cm. The soil temperature before sampling was listed in Table 1, with average temperature in Spring of 21.3°-22.0°C, Summer of 27.0°-28.0°C and Autumn of 18.0°-20.2°C. The air moisture tested in the same time was 82% for Spring, 86% for Summer, and 82.6% for Autumn. After collection, the soils from the same soil layer were mixed well in the field, sealed in bags, and immediately brought to the laboratory for measurement or stored in a −80°C freezer for soil microbial analysis. Other mixed soil samples were brought to the laboratory for water content test and air-dried for basic chemical properties analysis.

**Table 1.** Soil temperature and air moisture at different sampling stages in orchard

| Item | Spring | Summer | Autumn |
| --- | --- | --- | --- |
| Soil temperature (°C) |  |  |  |
| 0 cm | 21.3 | 27.5 | 18.0 |
| 5 cm | 22.0 | 27.7 | 19.0 |
| 10 cm | 21.5 | 28.0 | 19.5 |
| 15 cm | 21.7 | 27.3 | 20.2 |
| 20 cm | 21.4 | 27.0 | 19.7 |
| Air relative moisture (%) | 82.0 | 86.0 | 82.6 |

Soil temperature and air moisture was the average of two weeks data before sampling that was on April 27 for Spring, on August 22 for Summer and on November 4 for Autumn.

### 2.3. Chemical Analysis

Soil pH was measured in deionized water (1:5, soil:water). The total organic carbon (TOC) and total nitrogen (TN) were determined by potassium dichromate [20] and Kjeldahl digestion-distillation [21]. Exchangeable cations (K^+^, Na^+^, Mg^2+^ and Ca^2+^) were extracted by CH_3_COONH_4_ solution (pH = 7.0) [22], and analyzed using atomic absorption spectrophotometry (AA-6800, Shimadzu Corp., Kyoto, Japan). Soil samples was described in experimental section and conducted in triplicate.

### 2.4. Extraction and determination of microbial PLFAs

Four steps followed for soil PLFA extraction: 1). Five grams of soil was placed in a centrifuge tube. Then, 15 mL of a 0.2 M KOH (Sinopharm Chemical Reagent Co., Ltd., Shanghai, China) and methanol (Fisher Scientific Worldwide (Shanghai) Co., Ltd., Shanghai, China) solution was added before tightening the cap. The centrifuge tube was shaken for 5 min at 100 rpm. After shaking, the tube was incubated in a CU600 thermostat water bath for 5 min at 37°C. This procedure was repeated five times to help release the fatty acids from the soil sample. 2). Then, the tube was opened, and 3 mL of 1.0 M acetic acid (Sinopharm Chemical Reagent Co., Ltd) solution was added to decrease the pH of the reaction. 3). After adding 10 mL of n-hexane (Merck Co., Darmstadt, German) and mixing well, the tubes were centrifuged in an N00077 centrifuge for 15 min using the following settings: rotor: #12,150; speed: 2000 rpm; time: 15 min; and temperature: 4°C. After centrifugation, the supernatant n-hexane was transferred to a clean flask and air-dried under a fan. 4). The air-dried sample was re-suspended in 0.5 mL of a mixture of n-hexane:methyl-tert-butyl ether (Tedia Co., Inc., Fairfield, OH) (1:1, v/v) for 3-5 min and transferred to gas chromatography (GC) vials for PLFA determination.

Microbial PLFAs were determined using the Sherlock Microbial Identification System Sherlock MIS 4.5 (MIDI Inc., Newark, DE), including a 6890N Gas Chromatograph (GC) system (Agilent Technologies Inc., Palo Alto, CA), automatic injection devices, quartz capillary column, and flame ionization detector. The standard phospholipid fatty acid methyl ester (MIDI, Inc.) mixture and extracted samples were analyzed under the following chromatographic conditions: the temperature increment was controlled by the second-order program, with an initial temperature of 170°C that increased to 260°C at 5°C min^-1^ and then to 310°C at 40°C min^-1^ and maintained at 310°C for 90 sec; vaporization chamber temperature: 250°C; detector temperature: 300°C; carrier gas: H_2_ (2 mL min^-1^); blowing gas: N_2_ (30 mL min^-1^); pre-column pressure: 68.95 kPa; injection volume: 1 μL; injection split ratio: 100:1; and ionization mode: electron ionization (EI).

### 2.5. Calculation of Microbial Diversity

Known concentrations of 19:0 (nonadecanoic methyl ester) were added as internal standards and used to convert the retention-time peak areas to nanomoles per gram (nmol g^-1^) of soil (absolute abundance) and mole percent (mol %) (proportional abundance) of liquids. The absolute and proportional abundances of specific microbial groups were calculated by a summation of diagnostic lipid markers. The sum of 16:1 ω9c (PLFAs configuration type), 18:1 ω9c and 18:3 ω6c (6, 9, 12) was used to indicate fungi. The sum of 17:0 10 methyl and 18:0 10 methyl 18:1 ω9c and 18:3 ω6c (6, 9, 12) was used to indicate actinomycetes. The sum of 12:0, 14:0, 14:0 anteiso, 14:0 iso, 15:0 2OH, 15:0 3OH, 15:0 anteiso, 15:0 iso, 15:0 iso 3OH, 15:1 iso G, 16:0, 16:0 10 methyl, 16:0 2OH, 16:0 anteiso, 16:0 iso, 16:1 ω5c, 17:0 anteiso, 17:0 cyclo, 17:0 iso, 17:0 iso 3OH, 17:1 ω8c, 18:0, 18:0 iso, 18:1 ω7c, 18:1 ω7c 11 methyl, 18:3 ω6c (6,9,12), 19:0 cyclo ω8c, 19:0 iso, and 20:0 was used to indicate bacteria [23].

The number of species (Richness), Simpson diversity, Shannon-Wiener diversity and Alatalo evenness were used to calculate the ecological parameters of the microbial fatty acid biomarkers. The calculation equations are expressed as follows:

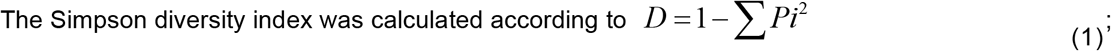

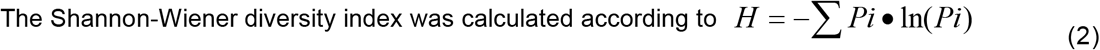

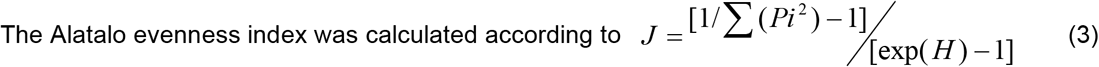

where 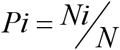; Ni is the content of the i^th^ kind of phospholipid fatty acid (PLFA); and N is the total PLFA content.

### 2.6. Statistical Analysis

An one-way ANOVA followed by least significant difference (LSD) multiple comparison test was used to establish significant differences among the means of soil properties, microbial community indicators (Bacterial PLFAs, Fungal PLFAs, Actinomycotic PLFAs, and B/F ratio of PLFAs), and microbial diversity indicators (Richness, Simpson index, Shannon-Wiener index and Alatalo index) in different seasons. A two-way analysis was used to establish significant differences in the total PLFA and individual PLFA variance between seasons and soil depths. A Spearman coefficient analysis was used to measure the correlation between soil properties and microbial indicators. A principle component analysis was used to measure the microbial community change between different seasons and soil depths. These analyses all used SPSS software version 17.0.

## Results

### 3.1. Soil Chemical Properties of Different Seasons in the Trial Orchard

The soils could be classified as clay, thermal and Typic Hapludult [24]. Principle chemical properties of the soil samples collected in the Spring, Summer and Autumn in the 0-5 and 5-20 cm depths are presented in Table 2.

**Table 2.** Chemical characteristics of soil sampling in different seasons at 0-5 and 5-20 cm soil depths in orchard

| Item | Spring | Summer | Autumn |
| --- | --- | --- | --- |
| 0-5 cm |  |  |  |
| pH | 4.87±0.06 a (5) | 4.46±0.08 b (5) | 4.95±0.04 a (5) |
| TOC (g kg<sup>-1</sup>) | 20.90±1.07 a | 12.32±1.53 b | 20.92±1.17 a |
| TN (g kg<sup>-1</sup>) | 1.46±0.24 bc | 1.14±0.08 c | 3.95±0.75 a |
| C/N ratio | 15.81±3.30 a | 10.76±0.93 b | 5.90±1.18 c |
| CEC (cmol(+) kg<sup>-1</sup>) | 9.42±0.26 a | 4.95±0.29 c | 7.09±0.25 b |
| Exchangeable K<sup>+</sup> (mg kg<sup>-1</sup>) | 124.00±15.64 a | 141.79±30.11 a | 99.67±13.16 b |
| Exchangeable Na<sup>+</sup> (mg kg<sup>-1</sup>) | 17.75±0.62 b | 30.67±3.61 a | 14.09±1.13 b |
| Exchangeable Mg<sup>2+</sup> (mg kg<sup>-1</sup>) | 30.14±3.83 a | 28.41±7.56 a | 24.80±2.79 a |
| Exchangeable Ca<sup>2+</sup> (mg kg<sup>-1</sup>) | 163.98±20.25 a | 122.72±20.74 a | 192.83±33.25 a |
| 5-20 cm |  |  |  |
| pH | 4.81±0.06 a (5) | 4.38±0.07 b (5) | 4.83±0.08 a (5) |
| TOC (g kg<sup>-1</sup>) | 19.15±0.92 a | 10.25±0.89 b | 18.78±0.67 a |
| TN (g kg<sup>-1</sup>) | 1.61±0.14 ab | 0.86±0.06 b | 2.20±0.25 a |
| C/N ratio | 12.85±0.81 ab | 11.88±0.55 b | 9.04±1.10 bc |
| CEC (cmol(+) kg<sup>-1</sup>) | 9.29±0.56 a | 5.34±0.39 c | 6.87±0.18 b |
| Exchangeable K<sup>+</sup> (mg kg<sup>-1</sup>) | 62.85±7.54 ab | 99.22±12.24 a | 55.95±7.85 ab |
| Exchangeable Na<sup>+</sup> (mg kg<sup>-1</sup>) | 15.22±1.26 b | 43.80±5.91 a | 10.21±0.85 b |
| Exchangeable Mg<sup>2+</sup> (mg kg<sup>-1</sup>) | 22.71±6.17 a | 34.47±8.80 a | 20.05±4.82 a |
| Exchangeable Ca<sup>2+</sup> (mg kg<sup>-1</sup>) | 121.38±22.12 a | 117.97±51.79 a | 164.97±18.31 a |

TOC = total organic carbon, TN = total nitrogen, CEC = cation exchange capacity, C/N ratio = ratio of total organic carbon to total nitrogen. Values followed by the same letter (s) in a low are not significantly different at $P < 0.05$ using LSD post hoc tests. Case number is shown in parentheses

Acidic soil pH (0-5 cm: 4.46; 5-20 cm: 4.38), total organic carbon (TOC) (0-5 cm: 12.32 g kg^-1^; 5-20 cm: 10.25 g kg^-1^), total nitrogen (TN) (0-5 cm: 1.14 g kg^-1^; 5-20 cm: 0.86 g kg^-1^), and cation-exchange capacity (CEC) (0-5 cm: 4.95 cmol(+) kg^-1^; 5-20 cm: 5.34 cmol(+) kg^-1^) were the lowest in the Summer samples, with significant differences (*P* < 0.05). However, the exchangeable K^+^ (0-5 cm: 141.79 g kg^-1^; 5-20 cm: 99.22 g kg^-1^) and Na+ (0-5 cm: 30.67 g kg^-1^; 5-20 cm: 43.80 g kg^-1^) were significantly higher than in the other seasons. Furthermore, the ratio of total organic carbon to total nitrogen (C/N ratio) in soil varied significantly between seasons, with the highest value in the Spring (0-5 cm: 15.81; 5-20 cm: 12.85), the middle value in the Summer (0-5 cm: 10.76; 5-20 cm: 11.88), and the lowest value in the Autumn (0-5 cm: 5.90; 5-20 cm: 9.04) (*P* < 0.05). No significant difference was detected between seasons in exchangeable Mg^2+^ or Ca^2+^ (*P* > 0.05).

No significant difference of pH, TOC, C/N ratio, CEC, exchangeable Mg^2+^ or Ca^2+^ was found between soil depths.

### 3.2. PLFAs of Total Microbial, Bacteria, Fungi and Actinomycotic of Different Seasons

The total microbial PLFAs peaked in the Summer in both the 0-5 and 5-20 cm soil depths (0-5 cm: 258.97 μg g^-1^ soil, *P* < 0.01; 5-20 cm: 270.99 μg g^-1^ soil, *P* < 0.01). The results of the two-way analysis demonstrated that the total microbial PLFAs were significantly different between seasons (*P* < 0.001). The quantities of bacterial, fungal and actinomycotic PLFAs in the orchard soil increased significantly in the Summer compared to those in the Spring and Autumn in the 0-5 and 5-20 cm soil depths (*P* < 0.01). In the 0-5 cm soil depth, the peak values of bacterial PLFAs, fungal PLFAs and actinomycotic PLFAs were 216.05, 33.94 and 8.96 μg g^-1^ soil, respectively. For the 5-20 cm soil depth, bacterial PLFAs, fungal PLFAs and actinomycotic PLFAs summit to 230.00, 31.15 and 9.83 μg g^-1^ soil, respectively (Table 3). Microbial PLFAs contents (μg g^-1^ soil) of different seasons sampled at 0-5 and 5-20 cm soil depths in orchard is shown in Table 4. Two-way analysis of variance for the effects of seasons (Spring, Summer, Autumn) and soil depths (0-5 cm, 5-20 cm) on microbial PLFAs contents are demonstrated on Table 5. Quantities of PLFAs for 29 microbe groups varied significantly between seasons, except for 15:0 iso 3OH, 15:1 iso G, 16:0 2OH, and 17:0 iso 3OH.

**Table 3.** Contents of total PLFAs, bacterial PLFAs, fungal PLFA, actinomycetical PLFAs, and the ratio of bacterial to fungal of soils sampling on different seasons at 0-5 and 5-20 cm depths in orchard

| Item | Spring | Summer | Autumn |
| --- | --- | --- | --- |
| 0-5 cm |  |  |  |
| Total PLFAs ($\mu\text{g g}^{-1}$ soil) | 14.07±8.23 c B (4) | 258.97±23.48 a A (5) | 99.93±18.62 b B (4) |
| Bacterial PLFAs ($\mu\text{g g}^{-1}$ soil) | 11.46±6.75 b B | 216.05±20.44 a A | 79.79±14.56 b B |
| Fungal PLFAs ($\mu\text{g g}^{-1}$ soil) | 2.27±1.23 c B | 33.94±3.96 a A | 16.09±3.64 b B |
| Actinomycetrical PLFAs ($\mu\text{g g}^{-1}$ soil) | 0.34±0.26 b B | 8.96±0.81 a A | 4.05±1.00 b B |
| B/F ratio | 4.79±0.29 a A | 6.37±0.61 a A | 5.15±0.60 a A |
| 5-20 cm |  |  |  |
| Total PLFAs ($\mu\text{g g}^{-1}$ soil) | 10.63±1.58 b B (5) | 270.99±58.94 a A (5) | 51.28±7.17 b B (5) |
| Bacterial PLFA ($\mu\text{g g}^{-1}$ soil) | 8.66±1.29 b B | 230.00±50.36 a A | 44.60±6.05 b B |
| Fungal PLFA ($\mu\text{g g}^{-1}$ soil) | 1.71±0.32 c B | 31.15±6.26 a A | 5.37±0.83 bc B |
| Actinomycetrical PLFA ($\mu\text{g g}^{-1}$ soil) | 0.24±0.05 b B | 9.83±2.45 a A | 1.30±0.60 b B |
| B/F ratio | 5.33±0.59 b B | 7.38±0.24 a AB | 8.70±1.07 a A |

Mean ± standard error. the values followed by the same lowercase and capital letter (s) in a column are not significantly different at $P < 0.05$ and $P < 0.01$ using LSD post hoc tests, respectively. Case number is shown in parentheses, 1 case was missed for Spring and Autumn at 0-5 cm depth respectively.

**Table 4.** Microbial PLFAs contents (μg g^-1^ soil) of different seasons sampled at 0-5 and 5-20 cm soil depths in orchard

| Fatty acid configuration | 0-5 cm |  |  | 5-20 cm |  |  |
| --- | --- | --- | --- | --- | --- | --- |
|  | Spring (4) | Summer (5) | Autumn (4) | Spring (5) | Summer (5) | Autumn (5) |
| 12:0 | 0.05±0.04 | 2.67±0.44 | 0.65±0.17 | 0.00±0.00 | 1.68±1.03 | 0.23±0.14 |
| 14:0 | 0.19±0.10 | 4.89±0.54 | 1.43±0.26 | 0.16±0.03 | 5.15±1.06 | 0.76±0.06 |
| 14:0 anteiso | 0.04±0.04 | 0.00±0.00 | 0.15±0.15 | 0.03±0.03 | 0.00±0.00 | 0.68±0.23 |
| 14:0 iso | 0.03±0.03 | 1.72±0.20 | 0.38±0.09 | 0.01±0.01 | 1.70±0.41 | 0.05±0.05 |
| 15:0 2OH | 0.06±0.06 | 0.37 ±0.18 | 0.00±0.00 | 0.01±0.01 | 0.46±0.19 | 0.00± 0.00 |
| 15:0 3OH | 0.19±0.14 | 3.99±0.88 | 1.14±0.37 | 0.18±0.04 | 4.33±0.86 | 0.61±0.30 |
| 15:0 anteiso | 0.40±0.15 | 9.07±1.17 | 3.22±0.68 | 0.43±0.08 | 9.25±1.85 | 2.13±0.23 |
| 15:0 iso | 0.58±0.38 | 15.08±0.91 | 5.44±1.15 | 0.22±0.07 | 13.97±5.74 | 2.74±0.86 |
| 15:0 iso 3OH | 0.02±0.02 | 0.00±0.00 | 0.00±0.00 | 0.07±0.04 | 0.00±0.00 | 0.00±0.00 |
| 15:1 iso G | 0.04±0.03 | 0.00±0.00 | 0.15±0.11 | 0.94±0.84 | 13.96±13.96 | 2.29±2.25 |
| 16:0 | 3.85±2.34 | 56.42±4.23 | 23.36±3.44 | 2.14±0.45 | 48.29±14.45 | 9.61±2.13 |
| 16:0 10 methyl | 0.68±0.36 | 16.04±1.90 | 5.92±1.22 | 0.47±0.07 | 17.58±3.72 | 3.54±0.82 |
| 16:0 2OH | 0.00±0.00 | 0.00±0.00 | 0.00±0.00 | 0.01±0.01 | 0.00±0.00 | 0.00±0.00 |
| 16:0 anteiso | 0.24±0.04 | 4.01±1.50 | 1.30±0.24 | 0.37±0.07 | 2.49±0.36 | 0.92±0.17 |
| 16:0 iso | 0.64±0.40 | 17.89±2.61 | 6.09±1.33 | 0.38±0.06 | 18.19±3.91 | 3.70±0.64 |
| 16:1 ω9c | 0.00±0.00 | 1.86±0.57 | 2.76±0.61 | 0.00±0.00 | 1.14±0.41 | 0.00±0.00 |
| 16:1 ω5c | 1.13±0.88 | 7.33±0.77 | 0.00±0.00 | 0.52±0.17 | 7.86±2.04 | 1.22±0.19 |
| 17:0 10 methyl | 0.14±0.11 | 3.22±0.26 | 1.33±0.28 | 0.08±0.02 | 3.89±0.76 | 0.59±0.20 |
| 17:0 anteiso | 0.46±0.14 | 8.94±1.36 | 3.06±0.64 | 0.52 ±0.11 | 8.70±1.52 | 1.78±0.17 |
| 17:0 cyclo | 0.09±0.07 | 4.38±0.77 | 1.04±0.29 | 0.08±0.01 | 4.31±0.77 | 0.52±0.15 |
| 17:0 iso | 0.48 ±0.25 | 17.13±2.33 | 5.35±1.21 | 0.33±0.05 | 18.53±3.27 | 3.42±0.28 |
| 17:0 iso 3OH | 0.00±0.00 | 0.00±0.00 | 0.00±0.00 | 0.12±0.07 | 0.00±0.00 | 0.00±0.00 |
| 17:1 ω8c | 0.08±0.08 | 2.23±0.49 | 0.40±0.18 | 0.02±0.02 | 1.73±0.50 | 0.00±0.00 |
| 18:0 | 0.70±0.35 | 12.99±1.20 | 5.89±1.07 | 0.66±0.13 | 13.39±2.69 | 3.19±0.37 |
| 18:0 iso | 0.03±0.03 | 2.24±0.24 | 0.47±0.27 | 0.03±0.02 | 2.73±0.52 | 0.20±0.13 |
| 18:0 10 methyl | 0.20±0.15 | 5.75±0.58 | 2.72±0.76 | 0.16±0.04 | 5.79±1.72 | 0.70±0.43 |
| 18:1 ω9c | 1.93±1.06 | 28.87±3.45 | 14.79±3.42 | 1.44±0.30 | 27.09±5.88 | 4.57±0.74 |
| 18:1 ω7c | 0.53±0.34 | 7.86±0.86 | 3.28±0.56 | 0.38±0.15 | 8.90±2.45 | 1.55±0.28 |
| 18:1 ω7c 11 methyl | 0.08±0.08 | 1.66±0.14 | 0.58±0.42 | 0.00±0.00 | 1.56±0.48 | 0.00±0.00 |
| 18:3 ω6c (6,9,12) | 0.33±0.17 | 3.21±0.80 | 1.30±0.23 | 0.28±0.03 | 3.06±0.40 | 0.79±0.24 |
| 19:0 cyclo ω8c | 0.65±0.38 | 15.95±0.72 | 6.39±1.43 | 0.43±0.06 | 22.14±5.22 | 4.83±0.84 |
| 19:0 iso | 0.01±0.01 | 0.53±0.32 | 0.00±0.00 | 0.01±0.01 | 0.44±0.14 | 0.00±0.00 |
| 20:0 | 0.20±0.10 | 2.67±0.10 | 1.34±0.24 | 0.16±0.04 | 2.66±0.62 | 0.63±0.12 |

Case number is shown in parentheses. Mean±standard error.

**Table 5.**
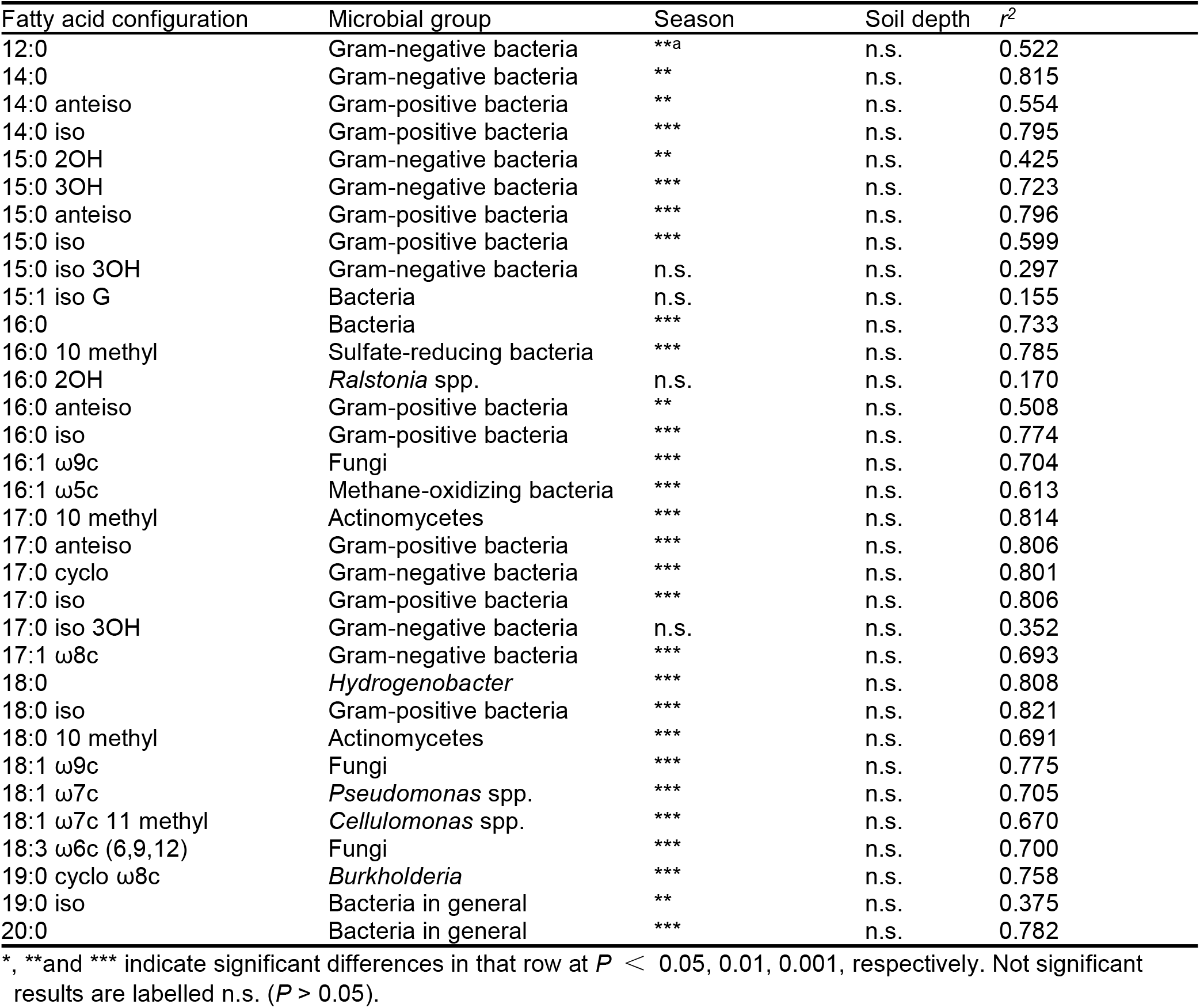
Two-way analysis of variance for the effects of seasons (Spring, Summer, Autumn) and soil depths (0-5 cm, 5-20 cm) on microbial PLFAs contents

### 3.3. Microbial Diversity Change between Seasons in the Trial Orchard

The microbial diversity analysis of the orchard soil showed that the Richness (number of microbial species) was in the range of 18 to 31, and the Simpson index, Shannon-Wiener index and Alatalo index were 0.76 to 0.93, 3.16 to 3.97 and 0.59 to 0.76, respectively (Fig. 2). The microbial diversities of the orchard soil in different seasons varied. In the 0-5 cm soil depth, the Richness and the Simpson, Shannon-Wiener and Alatalo indexes peaked in the Summer, with values of 30.0, 0.92, 3.91 and 0.65, respectively. The Simpson and Shannon-Wiener indexes in the Summer were significantly different from those in the Spring and Autumn (*P* < 0.05). A significantly difference in Richness was been found between the Summer and Spring. No significant difference in the Alatalo index between seasons was found (*P* > 0.05). In the 5-20 cm soil layer, the highest values of Richness and the Simpson and Shannon-Wiener index were still been found in the Summer, with values of 29.2, 0.92 and 3.81, respectively. The Richness and the Simpson index of the soil microbes in the Summer were significantly higher than those in the Spring (*P* < 0.05). There was no significant difference in the Shannon-Wiener index. However, the Alatalo index of the soil microbes in the Autumn (0.76) was significantly higher than that in the Spring (0.62) (*P* < 0.05). Overall, the microbial diversity of the orchard soil in the Summer was higher than in the Spring and Autumn.

**Fig. 2.**
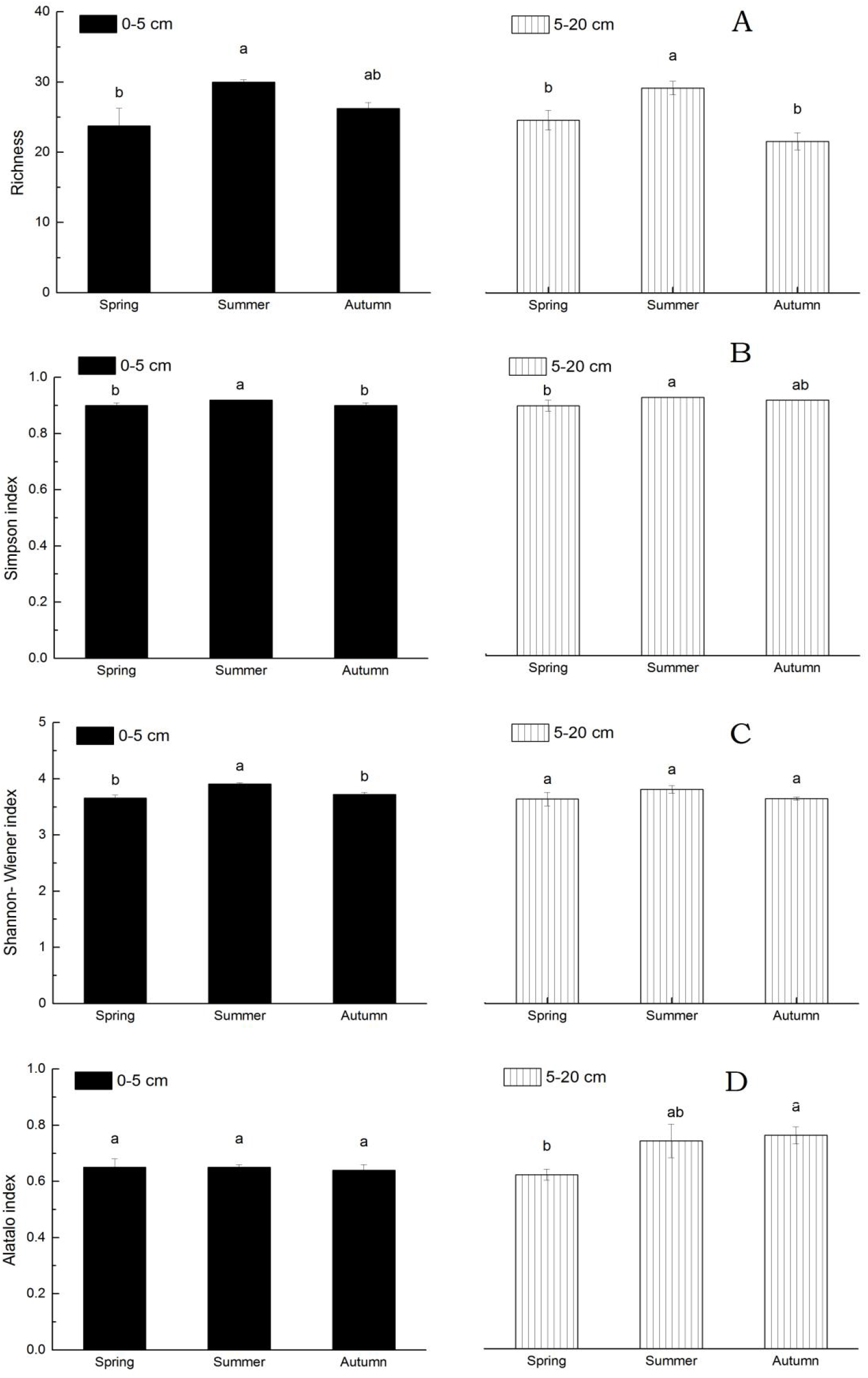
Microbial diversity of Richness (A) Simpson index (B), Shannon-Wiener index (C), and Alatalo index (D) in orchard soil of 0-5 and 5-20 cm s at different seasons. Bars with the same letter (s) are not significantly differences between seasons for each depth at *P* < 0.05 using LSD post hoc tests.

### 3.4. Correlation between Microbial Communities and Soil Properties

The correlation analysis was carried out between the soil properties (pH, TOC, TN, C/N ratio, CEC, and exchangeable K^+^, Na^+^, Mg^2+^ and Ca^2+^), microbial quantities (total PLFAs, bacterial PLFAs, fungal PLFAs, actinomycotic PLFAs, B/F ratio) and microbial diversities (Richness, the Simpson, Shannon-Wiener and Alatalo indexes). The results showed that the microbial quantities (total PLFAs, bacterial PLFAs, fungal PLFAs, and actinomycotic PLFAs) were significantly negatively correlated with the pH, TOC, TN and CEC, with Spearman correlation coefficients of-0.530 to −0.618 (*P* < 0.01), −0.572 to −0.642 (*P* < 0.01), −0.401 to −0.422 (*P* < 0.05) and −0.791 to −0.831 (*P* < 0.01), respectively (Table 5). The microbial diversity (Richness, and the Simpson and Shannon-Wiener indexes) was significantly negatively correlated with the pH, TOC, TN and CEC, with Spearman correlation coefficients of-0.460 to −0.753 (*P* < 0.05), −0.419 to −0.707 (*P* < 0.05), −0.450 to −0.526 (*P* < 0.05), and −0.446 to −0.722 (*P* < 0.05), respectively (Table 6).

**Table 6.** Spearman correlation coefficients matrix for microbial community variations and chemical properties in orchard soils

| Item | pH | TOC | TN | C/N<br>ratio | CEC | Exchangeable |  |  |  |
| --- | --- | --- | --- | --- | --- | --- | --- | --- | --- |
|  |  |  |  |  |  | K<sup>+</sup> | Na<sup>+</sup> | Mg<sup>2+</sup> | Ca<sup>2+</sup> |
| Total PLFAs | -0.614** | -0.617** | -0.401* | -0.302 | -0.817** | 0.284 | 0.493** | 0.119 | -0.177 |
| Bacterial PLFAs | -0.618** | -0.642** | -0.422* | -0.317 | -0.831** | 0.296 | 0.485** | 0.141 | -0.160 |
| Fungal PLFAs | -0.530** | -0.572** | -0.401* | -0.313 | -0.791** | 0.390* | 0.482** | 0.229 | -0.074 |
| Actinomycetical<br>PLFAs | -0.593** | -0.555** | -0.434* | -0.248 | -0.744** | 0.320 | 0.525** | 0.150 | -0.165 |
| B/F ratio | -0.517** | -0.565** | -0.354 | -0.119 | -0.595** | -0.259 | 0.131 | -0.189 | -0.263 |
| Richness | -0.460* | -0.419* | -0.526** | 0.051 | -0.446* | 0.403* | 0.622** | 0.332 | -0.085 |
| Simpson<br>diversity | -0.753** | -0.707** | -0.498** | -0.159 | -0.722** | 0.132 | 0.517** | -0.012 | -0.328 |
| Shanon-Wiener<br>diversity | -0.602** | -0.446* | -0.450* | 0.007 | -0.556** | 0.330 | 0.504** | 0.090 | -0.380* |
| Alatalo<br>evenness | -0.054 | 0.096 | 0.215 | -0.062 | 0.028 | -0.176 | -0.252 | -0.203 | -0.092 |

FLFA = phospholipid fatty acid, B/F ratio = ratio of bacterial to fungal PLFA, TOC = total organic carbon, TN = total nitrogen, CEC = cation exchange capacity, C/N ratio = ratio of total organic carbon to total nitrogen.
\*, \*\* indicate spearman coefficient significant differences at $P < 0.05, 0.01$ , respectively ( $n = 28$ ).

## Discussion

### 4.1. Microbial Population Variation of Soil Ecosystem Among Seasons

Usually, microbial population varies by moisture and temperature change among seasons. However, whether this change could be found easily depend on how long the suitable season last and how hard the extreme water and temperature condition affect on soil microorganisms. In subtropical mountain area, the soil was utilized as planting peach, obvious microbial change was confirmed in this article. And, principal component 1 explained 46.8 % of the variation in the soil microbial community (Fig. 3). The total, bacterial, fungal, and actinomycotic PLFAs, the B/F ratio, and the Richness were the main drivers of principal component 1 (Table S1). The remarkable change in microbial community was related to the peak soil microorganism growth in the Summer because of comfortable temperature and rainfall.

**Fig. 3.**
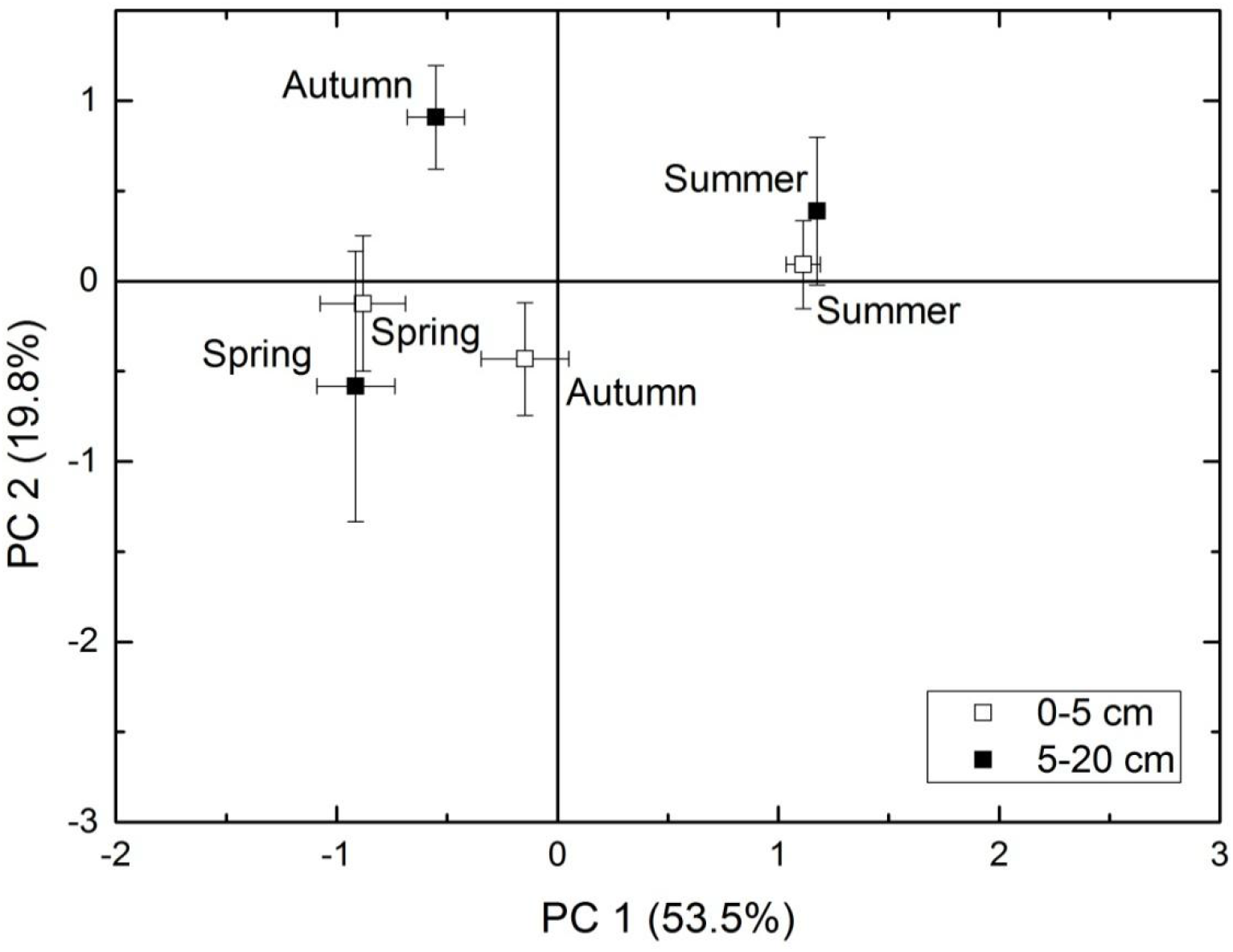
Principal components analysis (PCA) of microbial community from orchard soil sampled at different seasons and depths. Percent variance explained by each component (PC) is shown in parentheses. Error bars represent standard error (*n* = 28).

Principal component 2 explained 26.5 % of the variation in the soil microbial community (Fig. 3). The Simpson index and Shannon-Wiener index were its major drivers (Table S1). This could be explained by the microorganism propagating well and maintaining a good balance in the Summer in the tested orchard soil system.

The results of our study are consistent with the results of Zhu et al. [14] in an evergreen broadleaf forest and Qi et al. [15] in a bamboo grove in the subtropical climate zone. An investigation of an orchard [13], grassland [12] and forest [17] in the temperate monsoon climate zone was reported similar trends. However, unlike our results, Shi et al. [16] observed seasonal variations characterized by low values for most of the microbial biomass (C, N, and P), enzyme activities and PLFAs in the Summer in southwestern Quebec, Canada. This difference is related to the particularly dry climate (drier than the long-term average of the season) with a significantly lower moisture content than in the Spring and Autumn. It is well documented that temperature and moisture are the main factors related to microbial abundance and distribution.

### 4.2. Relationship between Soil Physicochemical Properties and Microbial Communities

The reported relationship between soil physicochemical properties and microbial communities varies among studies. In this study, in the view of distribution among seasons and soil depths, the total PLFAs, bacterial PLFAs, fungal PLFAs, actinomycotic PLFAs, number of species (Richness), Simpson and Shannon-Wiener diversity indexes were significantly negatively correlated with the soil TOC, TN and CEC. In the Summer, of most soil microbes experienced rapid growth, and a significant decrease in the soil pH, TOC, TN, and CEC was found (Table 1). Liu et al. [18] also found the microbial dominance index and Shannon-Wiener index to be negatively related to soil NH_4_^+^-N and NO_3_^-^-N in an apple system by observation on different growing periods. However, in the view of change by utilization, Yao et al. [6] illustrated microbial biomass C, basal respiration and total PLFA to be highly correlated with organic C and TN on red soil orchard ecosystem. In some ecosystem soil microbes grew rapidly with energy and nutrition consumption [25, 26]. It is clearly suggested that the propagation and growth of soil microorganisms from Spring to Summer require energy from TOC and nutrition from N in an subtropical orchard system. Microbes mainly act as “consumers” could be well documented.

## Conclusions

In subtropical orchards, the temperature and humidity in the Summer are conducive to the growth of soil microorganisms. PLFA analysis showed that the quantity of soil microbes in the soil samples collected in the Summer was significantly higher than in the Spring and Autumn. The Simpson and Shannon-Wiener indices also both peaked in the Summer sample collections. The total PLFAs, bacterial PLFAs, fungal PLFAs, actinomycotic PLFAs, Richness, and the Simpson and Shannon-Wiener indexes were significantly negatively correlated with seasonal changes in the soil pH, TOC, TN and CEC.

The function of microbial community in the translation and accumulation of soil nutrition from Summer to Autumn should be studied further. The changes in functional flora (including Archaea) in the soil microorganisms resulting from different orchard management strategies merit further investigations, and the relationships between the flora change and the orchard litter, soil organic matter (SOM), soil C/N ratio, ammonia N, nitrate N, pH, soil respiration, and other parameters should be merited further studied.

## Ackonwledgements

We thank Mr. Zheng Zhong for his support and long-term care of this stationary experiments.

## Author Contributions

**Funding acquisition:** BW XL.

**Project administration:** BW.

**Investigation:** XL.

**Chemical Analysis:** XL.

**Determination of microbial PLFAs:** GH.

**Statistical Analysis:** XL.

**Writing-original draft:** XL.

**Writing-review & editing:** MW BW.

## Supplemental Information

**Table S1.**
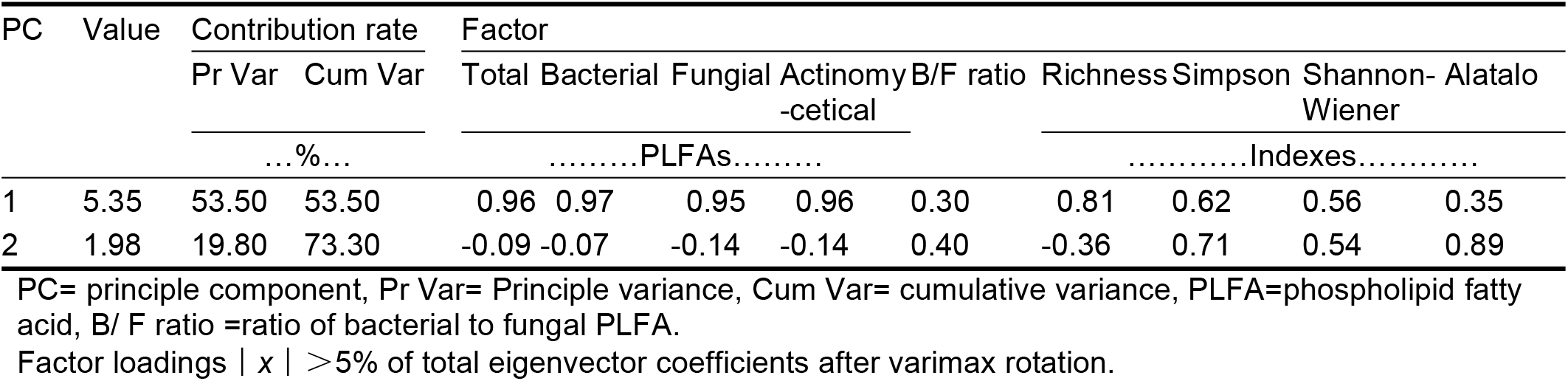
Principle component (PC) factors value and total eigenvector coefficients between PC factors and microbial community indexes after varimax rotation

## References

1. Simon S, Bouvier JC, Debras JF, Sauphanor B. Biodiversity and pest management in orchard systems. A review. Agron. Sustainable Dev. 2010; 30: 139–152.

2. Labrière N, Locatelli B, Laumonier Y, Freycon V, Bernoux M. Soil erosion in the humid tropics: A systematic quantitative review. Agric. Ecosyst. Environ. 2015; 203: 127–139.

3. Rowlings DW, Grace PR, Scheer C, Kiese R. Influence of nitrogen fertiliser application and timing on greenhouse gas emissions from a lychee *(Litchi chinensis)* orchard in humid subtropical Australia. Agric. Ecosyst. Environ. 2013; 179: 168–178.

4. Cai MF, Mcbride MB, Li KM. Bioaccessibility of Ba, Cu, Pb, and Zn in urban garden and orchard soils. Environ. Pollut. 2016; 208: 145–152.

5. Young IM, Crawford JW. Interactions and self-organization in the soil-microbe complex. Science (New series), 2004; 304: 1634–1637.

6. Yao H, He Z, Wilson MJ, Campbell CD. Microbial biomass and community structure in a sequence of soils with increasing fertility and changing land use. Microb. Ecol. 2000; 40: 223–237.

7. Li R, Khafipour E, Krause DO, Entz MH, de Kievit TR. Pyrosequencing reveals the influence of organic and conventional farming systems on bacterial communities. PLoS One; 2012; 7, e51897.

8. Mercier A, Dictor MC, Harris-Hellal J, Breeze D, Mouvet C. Distinct bacterial community structure of 3 tropical volcanic soils from banana plantations contaminated with chlordecone in Guadeloupe (French West Indies). Chemosphere. 2013; 92: 787–794.

9. Joa JH, Weon YH, Hyun HN, Jeun YC, Koh SW. Effect of long-term different fertilization on bacterial community structures and diversity in citrus orchard soil of volcanic ash. J. Microbiol. 2014; 52: 995–1001.

10. Yang DW, Zhang MK. Effects of land-use conversion from paddy field to orchard farm on soil microbial genetic diversity and community structure. Eur. J. Soil Biol. 2014; 64: 30–39.

11. Gilbert JA, Field D, Swift P, Thomas S, Cummings D, Temperton B, Weynberg K, Huse S, Hughes M, Joint I, Somerfield PJ, Mühling M, Rodriguez-Valera F. The taxonomic and functional diversity of microbes at a temperate voastal site: a ‘Multi-Omic’ study of seasonal and diel temporal variation. PloS One. 2010; 5: 1–17.

12. Bardgett RD, Lovell RD, Hobbs PJ, Jarvis SC. Seasonal changes in soil microbial communities along a fertility gradient of temperate grasslands. Soil Biol. Biochem. 1999; 31: 1021–1030.

13. Shishido M, Sakamoto K, Yokoyama H, Momma N, Miyashita S. Changes in microbial communities in an apple orchard and its adjacent bush soil in response to season, land-use, and violet root rot infestation. Soil Biol. Biochem. 2008; 40: 1460–1473.

14. Zhu WZ, Cai XH, Liu XL, Wang JX, Cheng JX, Cheng S, Zhang XY, Li DY, Li MH. Soil microbial population dynamics along a chronosequence of moist evergreen broad-leaved forest succession in southwestern China. J. Mount. Sci. 2010; 7: 327–338.

15. Qi LH, Ai WS, Fan SH, Du MY, Meng Y, Mao C. Soil microbial biomass carbon dynamics of *Phyllostachys edulis* forests under different managing patterns in the hilly region of central Hunan, southern China. J. Nanjing Forest Univ. 2013; 37: 45–48 (in Chinese with English abstract).

16. Shi Y, Lalande R, Hamel C, Ziadi N, Gagnon B, Hu Z. Seasonal variation of microbial biomass, activity, and community structure in soil under different tillage and phosphorus management practices. Biol. Fert. Soils. 2013; 49: 803–818.

17. Kim CS, Nam JW, Jo JW, Kim SY, Han JG, Hyun MW, Sung GH, Han SK. Studies on seasonal dynamics of soil-higher fungal communities in Mongolian oak-dominant Gwangneung forest in Korea. J. Microbiol. 2016; 54:14–22.

18. Liu LZ, Qin SJ, Lu DG, Wang BY, Yang ZY. Variation of potential nitrification and ammonia-oxidizing bacterial community with plant-growing period in apple orchard soil. J. Integr. Agric. 2014; 13: 415–425.

19. Zhuan WM. Map of Fujian Province. Fuzhou: Fujian provincial map and atlas publishing house; 2008.

20. Nelson DW, Sommers LE. Total carbon, organic carbon and organic matter. In Page AL, Miller RH. and Keeney DR. editors. Methods of Soil Analysis. Madison WI: Soil Sci. Soc. Am. 1982; 539–577.

21. Bremner JM, Mulvaney CS. Total nitrogen. In: Page AL, Miller RH, Keeney DR, editors. Methods of Soil Analysis. Madison WI: Soil Sci. Soc. Am.; 1982.

22. Jackson ML. Soil Chemical Analysis. Published by Author, 2nd ed. Madison WI; 1979.

23. Zhu YJ, Hu GP, Liu B, Xie HA, Zheng XF, Zhang JF. Using phospholipid fatty acid technique to analysis the rhizosphere specific microbial community of seven hybrid rice cultivars. J. Integr. Agric. 2012; 11: 1817–1827.

24. Soil Survey Staff. Keys to Soil Taxonomy. 11th ed. Washington, DC: USDA-Natural Resources Conservation Service; 2010.

25. Huang PM, Wang MK, Chiu CY. Soil mineral-organic matter-microbe interactions: impacts on biogeochemical processes and biodiversity in soils. Pedobiogia. 2005; 49: 609–635.

26. Huang PM, Wang SL, Tzou YM, Huang YB, Weng BQ, Zhuang SY, Wang MK. Physicochemical and biological interfacial interactions: impacts on soil ecosystem and biodiversity. Environ. Earth. Sci. 2013; 28: 2199–2209.

